# Indel-driven evolution of the canavanine tRNA-editing deacetylase enzyme CtdA

**DOI:** 10.1101/2024.11.30.626143

**Authors:** Nino Tabagari, Franziskus Hauth, Jennifer R. Fleming, Jörg S. Hartig, Olga Mayans

**Author notes:** These authors contributed equally (Nino Tabagari, Franziskus Hauth).

## Abstract

Proteins are heteropolymers composed of twenty standard amino acids. However, over 500 non-proteogenic amino acids exist in nature that can get misincorporated into proteins. Canavanine is an antimetabolite of L-arginine, with which it shares high chemical similarity. It can be utilized by bacteria such as *Pseudomonas canavaninivorans* in the legume rhizome as a sole source of carbon and nitrogen. However, canavanine is also incorporated in proteins of this bacterium as its arginyl-tRNA synthetase loads tRNA^Arg^ with both canavanine and arginine. The recently discovered canavanyl-tRNA^Arg^ deacetylase (CtdA) removes canavanine from misloaded tRNA^Arg^ and thereby prevents its incorporation in proteins. CtdA is the first enzyme known to edit tRNA mischarged with a non-proteinogenic amino acid. We have elucidated its crystal structure to 1.5 Å resolution and studied its active site using site-directed mutagenesis. We found that CtdA is a small monomeric enzyme that presents a central, deep cavity that predictably constitutes the canavanine binding site and a positively charged surface area that likely coordinates the CCA-3’ tRNA attachment sequence. The stand-alone, trans-editing CtdA is distantly related to the B3/B4 cis-editing domains of the large multi-subunit enzyme Phenylalanine tRNA synthetase (PheRS). Our comparative study reveals that CdtA and B3/B4 domains from bacterial and archeal/eukaryotic origin are three subclasses of a same conserved 3D-fold that differ in type-specific indels, which distinctly shape the substrate binding cleft of these proteins. We propose a unifying nomenclature of secondary structure elements for this 3D-fold. In CtdA, residues E191, Y104, N105 and E118 prove to be relevant for catalysis, of which N105 is conserved in bacterial B3/B4 domains. No other shared residues of catalytic relevance could be identified across enzymes of this class, so that a shared mechanism of catalysis appears unlikely in these editing enzymes.

## Introduction

Ensuring translational fidelity is a crucial task in all domains of life as misincorporation of incorrect amino acids into proteins can drastically reduce organismal fitness (Bacher et al., 2005; Mohler et al., 2017). Generally, faulty proteins reduce fitness through three phenomena: (i) loss of protein function, (ii) gain of protein toxicity and (iii) clean-up costs of erroneous proteins that can overload the turn-over machinery of the cell (Drummond & Wilke, 2009). Translational mistakes can either happen at the ribosome during decoding and polypeptide formation or earlier, during amino acid activation and tRNA charging. The latter process is mediated by aminoacyl-tRNA-synthetase (aaRS) enzymes. Typically, aaRSs are specific for the charging of a single proteinogenic amino acid onto its cognate tRNA, which is remarkable considering the vast number of other proteinogenic and non-proteinogenic amino acids that exist in the cell (Fichtner et al., 2017). To ensure translational accuracy, aaRS enzymes have developed a so-called ‘double-sieve’ mechanism that ensures correct amino acid charging. Here, a first active site within the enzyme activates and transfers the correct amino acid onto its tRNA, while a second, editing active site does proofreading and hydrolyzes incorrectly loaded aminoacyl-tRNAs (Rubio Gomez & Ibba, 2020; Jani & Pappachan, 2022). Approximately half of all aaRSs are known to rely on such *cis*-editing to efficiently load tRNA with cognate amino acids instead of structurally similar, non-cognate ones (Rubio Gomez & Ibba, 2020). In addition, a third ‘sieve’ mechanism has been discovered in both prokaryotes and eukaryotes that is performed by autonomous editing enzymes, which catalyze the in-*trans* editing of mischarged tRNAs (Jani & Pappachan, 2022).

A total of 23 aaRSs are known to date, one for each of the 20 proteinogenic amino acids (except for lysine, for which there are two) plus pyrrolysyl-tRNA synthetase (PylRS) and phosphoseryl-tRNA synthetase (SepRS) (Rubio Gomez & Ibba, 2020). In contrast, the number of freestanding trans-editing enzymes identified to date is more limited, currently including Ser-tRNA^Thr^ (which edits Ser loaded onto a Thr specific tRNA), Ser/Gly-tRNA^Ala^ (AlaX), D-aa-tRNA^aa^ (DTD), Cys-tRNA^Pro^ (Ybak), Ala/Cys-tRNA^Pro^ (ProXp-X) and other less specific ProXp variants that deacylate a variety of mischarged tRNAs (Jani & Pappachan, 2022). At exception of DTD, standalone trans-editing enzymes share sequence conservation with the cis-editing domains of the corresponding aaRS enzyme whose mischarged amino acids they correct for; namely, Ser-tRNA^Thr^ relates to the N2 domain of ThrRS, the AlaX trans-editing enzymes relate to the C-terminal domain of AlaRS, and Ybak and ProXp enzymes relate to the INS domain of ProRS (Jani & Pappachan, 2022). Recently, we have identified a canavanyl-tRNA^Arg^ deacetylase (CtdA) as a standalone editing enzyme that constitutes the first example of trans-editing enzyme with similarity to the *cis*-editing B3/B4 domain of the large beta subunit of Phenylalanine-tRNA-synthetases (PheRS), which corrects for mischarged tyrosyl-tRNA^Phe^ (Hauth et al., 2023). This relation is unexpected as CtdA specifically hydrolyzes the non-proteinogenic arginine antimetabolite canavanine from tRNA^Arg^, which differs notably from the phenyl-/tyrosyl-substrates processed by PheRS. Canavanine (δ-oxa-arginine) and arginine differ only in a single atom substitution (atom δO instead of δC) in the lateral side chain. CtdA is found in bacteria that encounter canavanine in their natural habitat, often originating from leguminous plants. The enzyme enables the bacteria to circumvent canavanine intoxication caused by the misincorporation of this non-proteinaceous aminoacid into proteins in lieu of arginine. Thus, CtdA is also the first example to date of a non-proteinogenic editing enzyme.

PheRS is the aaRS enzyme best characterized at the molecular level so far. Three types of PhRS enzymes are currently known: bacterial (αβ)2-PheRS, eukaryotic/archaeal cytosolic (αβ)2-PheRS, and mitochondrial α-PheRS (α and β referring here to subunit composition). Three-dimensional structures of PheRS representatives from these three classes are available: namely, the cytosolic (Finarov et al., 2010) and mitochondrial enzymes (Klipcan et al., 2012) from *Homo sapiens* and the cytosolic PheRS from *Plasmodium vivax* (Sharma et al., 2021) among eukaryotes; PheRS from *Escherichia coli* and *Acinetobacter baumannii* (Baidin et al., 2021) and *Mycobacterium tuberculosis* (Michalska et al., 2021) among bacteria; and the enzyme from *Thermus thermophilus* (Moor et al., 2006) in archaea. In addition, an N-terminal fragment of the beta subunit of PheRS from the archaeon *Pyrococcus horikoshii* has also been characterized structurally (Sasaki et al., 2006). PheRS enzymes diverge significantly in primary sequence, domain composition and subunit organization, but crystal structures have revealed that their editing B3/B4 domains share a similar fold that carries sequence insertions characteristic of every PheRS type (Sasaki et al., 2006), where the editing pocket of the archaeal/eukaryotic domains differs drastically from that of their bacterial counterparts at the sequence and structure level. This divergence has made it difficult to establish whether the same catalytic mechanism is shared by these *cis*-editing domains.

In this work, we have structurally characterized CtdA, a distant homologue of the B3/B4 domain class that discriminates between the proteinogenic and non-proteinogenic amino acids arginine and canavanine, respectively. Specifically, we have solved the crystal structure of CtdA from *Pseudomonas canavaninivorans*, modelled the binding of the substrate canavanine, tested single mutation variants of CtdA to identify residues relevant for substrate binding or catalysis, and performed sequence and structure comparisons of CtdA and homologous B3/B4 editing domains from PheRS enzymes. Our findings suggest that the folds of CtdA and B3/B4 domains from PheRS share a closely conserved central scaffold, but have diverged evolutionarily via indels that define the topography of their substrate binding sites. xCommon catalytic residues across CtdA and B3/B4 domains could not be detected, so that the possible existence of a shared mechanism of catalysis remains unclear.

## Results

### CtdA is a small monomeric enzyme with an α/β fold

The atomic structure of CtdA from *Pseudomonas canavaninivorans* has been elucidated to 1.50 Å resolution using X-ray crystallography (**Fig 1A**, **Table 1**). Two essentially identical molecular copies of CtdA (RMSD_Cα_=0.32 Å for all 231 Cα atoms in the protein chain) are present in the asymmetric unit of the crystal form obtained in this study. An analysis of the molecular interface in the crystallographic dimer using PISA (Krissinel & Henrick, 2007; https://www.ebi.ac.uk/pdbe/pisa) suggested the dimer not to be physiological, but resulting from packing in the crystal lattice. This conclusion was supported by data from size-exclusion chromatography that indicated the presence in solution of a protein species with a molecular mass of ∼25.2 kDa (**Supplementary Fig S1**). This agrees excellently with the molecular mass of CtdA calculated from its amino acid sequence (25.0 kDa). Taken together, this data suggested that CtdA is monomeric in solution.

**Figure 1:**
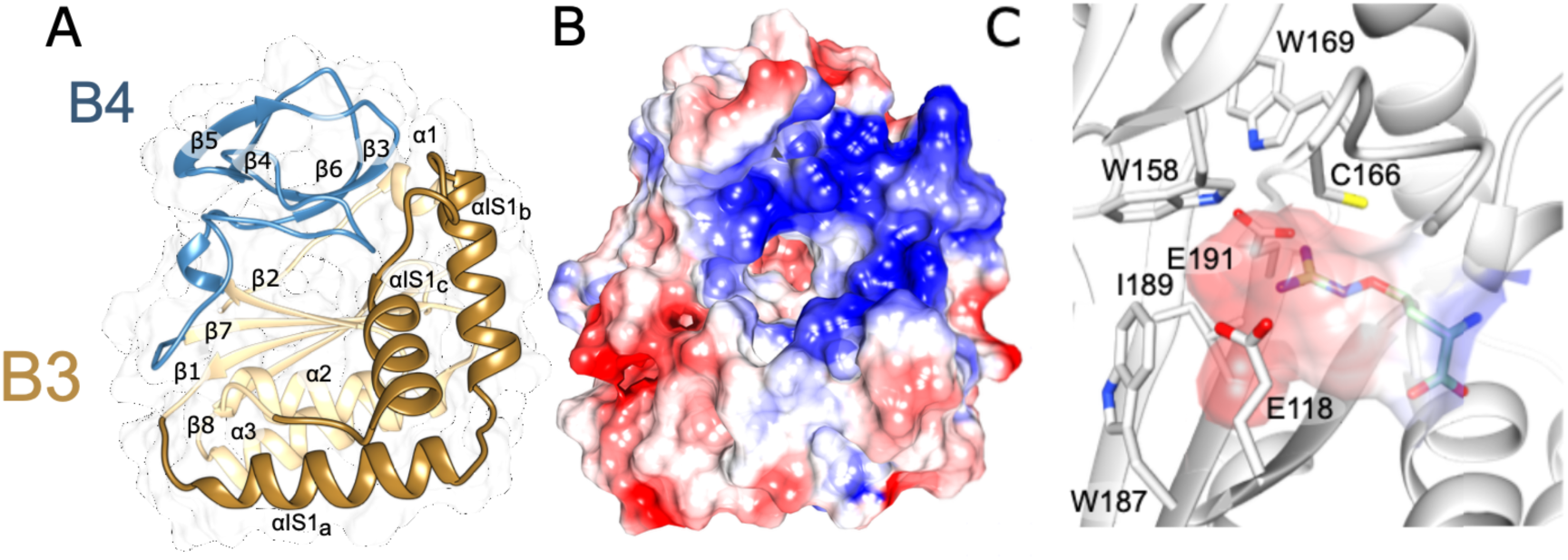
Crystal structure of CtdA. **A.** 3D-structure of CtdA coloured according to its homology to the B3 (gold; residues 1-120 and 184-231) and B4 (blue; residues 121-183) domains of PheRS. Within the B3 homology domains, dark gold indicates the long indel sequence (residues 31-95) unique to CtdA; **B.** Surface representation of CtdA coloured according to Coulomb’s electrostatic potential (red: -10 kcal/mol*e, blue: 10 kcal/mol*e). A central, deep cavity surrounded by positive electrostatic potential is here predicted to constitute the substrate binding site of CtdA; **C.** Close-up of the central cavity predicted to constitute the canavanine binding site of CtdA. Enzyme residues lining the central cavity are shown and annotated. The cavity is represented by its electrostatic surface. A canavanine molecule, docked using AutoDock (Trott et al., 2010), is depicted. All groups are coloured according to atomic composition, where: C, light gray; N, blue; O, red; S, yellow. To ease identification, in canavanine, C atoms are coloured green.

**Table 1:**
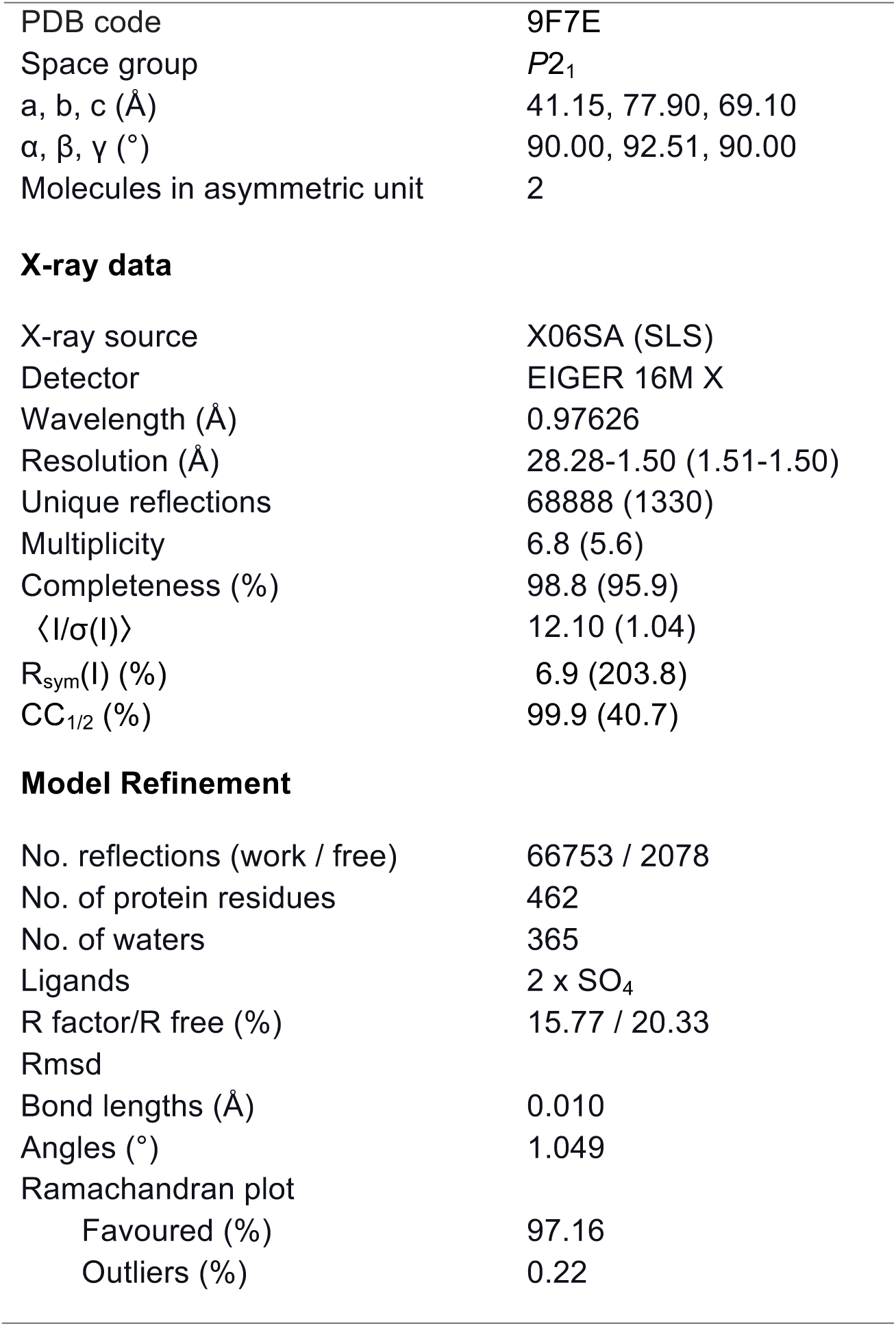
CtdA X-ray data statistics and model parameters.

|  |  |
| --- | --- |
| PDB code | 9F7E |
| Space group | $P2_1$ |
| a, b, c (Å) | 41.15, 77.90, 69.10 |
| $\alpha$, $\beta$, $\gamma$ (°) | 90.00, 92.51, 90.00 |
| Molecules in asymmetric unit | 2 |
| **X-ray data** |  |
| X-ray source | X06SA (SLS) |
| Detector | EIGER 16M X |
| Wavelength (Å) | 0.97626 |
| Resolution (Å) | 28.28-1.50 (1.51-1.50) |
| Unique reflections | 68888 (1330) |
| Multiplicity | 6.8 (5.6) |
| Completeness (%) | 98.8 (95.9) |
| $\langle I/\sigma(I) \rangle$ | 12.10 (1.04) |
| $R_{\text{sym}}(I)$ (%) | 6.9 (203.8) |
| $CC_{1/2}$ (%) | 99.9 (40.7) |
| **Model Refinement** |  |
| No. reflections (work / free) | 66753 / 2078 |
| No. of protein residues | 462 |
| No. of waters | 365 |
| Ligands | 2 x $\text{SO}_4$ |
| R factor/R free (%) | 15.77 / 20.33 |
| Rmsd |  |
| Bond lengths (Å) | 0.010 |
| Angles (°) | 1.049 |
| Ramachandran plot |  |
| Favoured (%) | 97.16 |
| Outliers (%) | 0.22 |

The 3D-fold of CtdA consists of seven alpha helices and eight beta strands that fold into a central four-stranded antiparallel β-sheet (formed by strands β1, β2, β7 and β8), which is capped by an apical β-rich domain at one side and packs against a bundle of three α-helices (α2, α3 and αIS1a) on its other face. An α-hairpin-like structure (formed by helices αIS1b and αIS1c) packs orthogonally against the helical bundle and central β-sheet (**Fig 1A**) (the nomenclature of secondary structure elements is described later below). The overall fold resembles that of domains B3/B4 in PheRS, with the apical β-rich domain sharing structural similarity with domain B4 and the central β-sheet and its flanking α-helical bundle with domain B3 (**Fig 1A**).

### Identification of the substrate binding site in CtdA

To identify the substrate binding site in CtdA, we performed an analysis of its surface topography. This revealed a deep cavity on the protein surface, centrally located in the fold, whose entrance is surrounded by a number of positively charged residues (K73, R76, K86, R87, R90, R167, R168, R172, R178) (**Fig 1B**). The cavity itself is largely lined up by hydrophobic residues (W158, W169, W187, I189) and contains negative charges at its deepest end (contributed by residues E118 and E191) (**Fig 1C**). Predictably, it accommodates the canavanyl-moiety of the substrate. The positively charged surrounding of the cavity likely mediates the binding of the negatively charged tRNA backbone, specifically the CCA-3’ sequence. This sequence is conserved at the 3’ end of all mature tRNA molecules and functions as the site of amino acid attachment. This sequence is important for the recognition of tRNA by enzymes. Large, multi-subunit PheRS enzymes have a large substrate binding interface and provide many opportunities for determinants of substrate specificity, so that their B3/B4 editing domains are not highly dependent on reading the tRNA sequence directly. On the contrary, in the small standalone CtdA, recognition and reading of the tRNA moiety is predictably critical to establish that the non-proteinogenic canavanine has been wrongly attached to tRNA.

In order to confirm that the identified cavity could accommodate the aminoacyl rest of the liganded tRNA and to get insights into essential substrate-binding amino acids, we performed co-crystallization as well as crystal soaking trials of CtdA with canavanine, arginine, *in vitro* transcribed tRNA^Arg^ or a synthetic RNA hairpin loop mimicking the tRNA CCA-3’ sequence. Unfortunately, these efforts were unsuccessful and a liganded CtdA structure could not be obtained. Thus, we performed instead *in silico* ligand docking studies of canavanine into the identified surface cavity using AutoDock (Trott et al., 2010) (**Fig 1C**). The resulting model revealed a canavanine molecule in extended conformation bound into the cavity, confirming that the dimensions of the cavity are suitable to accommodate the canavanyl moiety of the canavanyl-tRNA^Arg^ substrate. It further suggested that the negatively charged glutamate residues at the deep end of the cavity likely coordinate the positively charged guanidinium group of canavanine. Interestingly, the model also indicated that residue C166 might interact with atom δO in canavanine, thereby potentially playing a role in the discrimination between the highly similar canavanyl-and arginyl-substrates by the enzyme. To bring support to the hypothesis that CtdA binds differently to these substrates, we attempted to measure experimentally CtdA’s affinity for canavanine and arginine. However, our efforts to quantitate the binding of these ligands to CtdA using microscale thermophoresis were unsuccessful. The result led support to the view that CtdA has low affinity for the free amino acids and that engagement of the tRNA moiety by the cluster of positively charged residues must play an important role in the productive binding of the substrate.

Building on these structural and bioinformatics results, we aimed next to confirm functionally relevant residues playing a role in catalysis or substrate binding. For this, we first sought and investigated CtdA-related sequences to confirm the conservation of residues lining the predicted canavanyl-binding cavity in CtdA or areas around the cavity potentially co-localizing with the canavanyl-ribose ester bond to be excised. A sequence similarity search using BLAST (https://blast.ncbi.nlm.nih.gov) revealed 7 sequences with high similarity (∼66-30% seq. id.) to CtdA from *P. canavaninivorans* in this study (**Supplementary Fig S2**). The high similarity suggests that also those proteins might display canavanyl-tRNA^Arg^ deacylase activity. All sequences identified were from bacterial organisms, including pathogens such as *M. tuberculosis*, suggesting that this activity might be more widely spread among bacteria present in soil or animal digestive tracts (environments where legumes are present) than previously anticipated. In the sequences of those putative CtdA enzymes, residues suggested by crystal structures to be of significance to substrate binding (E118, C166, E191) (**Fig 2A**) or predicted to colocalize with the canavanyl-ribose ester bond in the substrate bound state (Y104, N105, S108; **Fig 2A**) were strictly conserved (**Supplementary Fig S2**). As strict conservation might point to functional significance, we set to test the contribution of these residues to catalysis. For this, we used site directed mutagenesis to generate single point mutants of CtdA for the following residue exchanges (designed based on structural tolerance predicted using FoldX; Schymkowitz et al., 2005) (**Supplementary Fig S3**): S80A, Y104F, N105D, S108A, E118Q, C166G, E191V. We then tested the performance of the CtdA mutated variants in canavanyl-tRNA^Arg^ editing *in vitro* (**Fig 2B**). To assess editing activity, tRNA^Arg^ was first canavanylated using an arginyl-tRNA^Arg^-synthesase as described previously (Hauth et al., 2023). Then, canavanyl-tRNA^Arg^ was incubated with the mutated CtdA variants. Various residue exchanges impacted editing activity strongly. For the exchanges E191V, Y104F, N105D, E118Q, after a 10 minute incubation time the percentage of canavanylated tRNA within the samples was similar to the no enzyme control samples, indicating that the mutation had abolished CtdA catalysis (**Fig 2B**). Mutants C166G, S080A and S108A still yielded editing activity, with S080A showing no difference to the wild-type enzyme. This result suggested that the negative charges at the deep end of the binding cavity, E191 and E118, are individually required for successful substrate binding and/or catalysis and that, similarly, residues N105 and Y104 predictably in the proximity of the canavanly-ribose junction are catalytically important. Conceivably, residues Y104 and N105 might form part of the catalytic oxyanion hole known to be required for the stabilization of the catalytic intermediate in hydrolase-mediated reactions (Bauer et al., 2020). Residue G117, which binds the hydroxyl group of canavanine in the docked model and is part of the sequence VGGEN, could also be part of the oxyanion hole. The participation of Y104 and/or G117 in oxyanion formation in CtdA would be in agreement with known oxyanion hole signature motifs (GX-, GGGX-and Y-motifs) in α/β-hydrolases (Bauer et al., 2020). On the contrary, activity data show that residue C166 does not appear to play a deterministic role in the binding or catalysis of canavanyl-tRNA^Arg^. Therefore, its significance in the discriminatory catalysis of canavanyl-versus arginyl-ester hydrolysis by CtdA remains uncertain.

**Fig 2:**
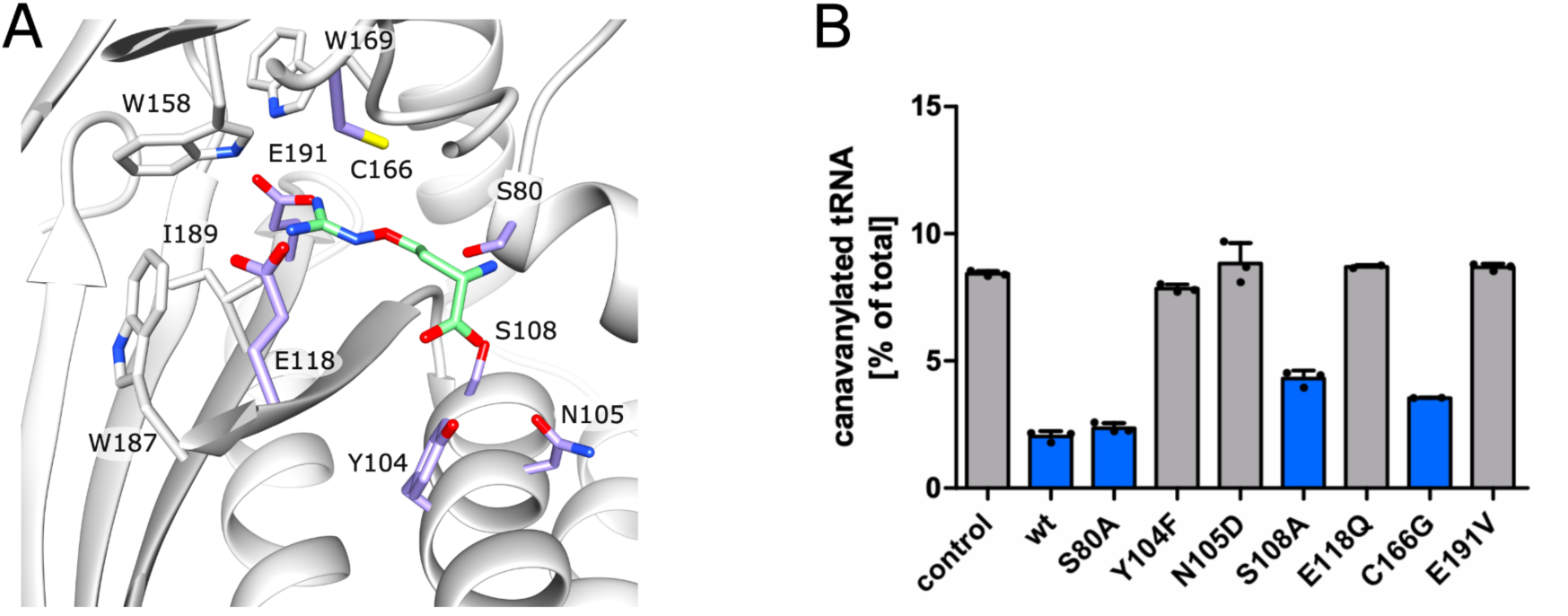
Site-directed mutagenesis studies of active site candidate residues in CtdA. **A.** Candidate active site residues in CtdA and their location relative to the predicted binding site for the canavanyl-rest (canavanine is shown in green; details as in Fig 1C). Residues mutated in this study are coloured lilac; **B.** Editing activity of CtdA and its mutated variants expressed as the percentage of canavanylated tRNA after 10 minutes incubation with the respective enzyme (n=3). A detailed description of the method is given in the Materials section. Original experimental data are shown in **Supplementary Fig S4**. Residue exchanges coloured in blue did not impair CtdA editing behaviour.

### The CtdA fold has evolved via indels

Next, we aimed to understand the relation of CtdA and B3/B4 domain homologues. For this, we first performed a structural search of available 3D-structures using DALI (Holm, 2022), which confirmed that CtdA shares close structural similarity with the β-subunit of PheRS enzymes. The result showed that CtdA relates to (in order of similarity) the B3/B4 domains of PheRS from *Pseudomonas aeuriginosa* (PDB entry 4P71; Abibi et al., 2014), *Eschericha coli* (3PCO; Mermershtain et al., 2011), *Acinetobacter baumannii* (6P8T; Baidin et al., 2021), *Staphylococcus haemolyticus* (2RHQ; Evdokimov et al., 2008), *Mycobacterium tuberculosis* (7DB7; Wang et al., 2021), *Pyrococcus horikoshii* (2CXI; Sasaki et al., 2006), *Thermus thermophilus* (2IY5; Moor et al., 2006), *Plasmodium vivax* (7BY6; Sharma et al., 2021), and *Homo sapiens* (3L4G; Finarov et al., 2010). Thus, CtdA shows a modestly closer similarity to the bacterial versus the archeal/eukaryotic B3/B4 domain types.

Following, we performed a structural comparison of CtdA with the B3/B4 homologues so-identified. The comparison revealed that CtdA and B3/B4 domains share a common structural core scaffold (**Fig 3A**), but that three indels at three different topological positions in the fold determine fold subtypes (**Fig 3B**). Indels in B3/B4 domains had been previously identified (Sasaki et al., 2006). By defining a unifying nomenclature for this enzyme class, where secondary structure elements within the shared core fold are numbered sequentially and the prefix IS is used to refer to elements in insertion sequences, we found that the first indel (insertion sequence 1; IS1) occurs between strand β1 and helix α2, the second indel (insertion sequence 2; IS2) between strands β5 and β6, and the third indel (insertion sequence 3; IS3) between strand β7 and helix α3 (**Fig 3C & Fig 4**). CtdA contains a single indel, IS1, while bacterial B3/B4 domains contain IS1 and IS3, and the archeal/eukaryotic type has IS2 (in this type, a short, single helical turn sequence that constitutes an N-terminal extension to helix α2 part of the common fold is found in IS1). In CtdA, IS1 is a 64-residue long sequence (notably longer than that of bacterial B3/B4 domains) that folds into three α-helices (helices α-IS1a, α-IS1b, α-IS1c). These form an L-shaped structure, where helix α-I1Sa lies perpendicular to helices α-IS1b and α-IS1c that form an α-hairpin (**Fig 1A, 3C**). In bacteria B3/B4 domains, IS1 is short and folds into a single α-helix (α-IS1a) that is structurally equivalent to that of CtdA (**Fig 3C**). Remarkably, both indels IS2 in archeal/eukaryotic B3/B4 domains and IS3 in bacterial B3/B4 domains fold into an α-hairpin that, although contributed by different topological parts of the protein chain, localize to the same topographical region of the 3D-fold (**Fig 3C**). This locus of the fold is approximately coincident with that occupied by the α-hairpin in indel IS1 in CtdA. Hence, in all cases indels fold against the shared fold scaffold and the same region of the surface topography of these enzymes is sculpted by indels -specifically, this is the surface surrounding the substrate binding site. In CtdA, the indel sequence is rich in positively charged residues and forms the entry to the aminoacyl-binding cavity, predictably playing a role in coordinating the CCA-3’ ribonucleotide moiety of the substrate. In the B3/B4 domain from *T. thermophilus,* IS3 also contains positively charged residues that interact with tRNA (Moor et al., 2006). However, in B3/B4 domains features indicative of tRNA binding are less prominent (**Fig 3C**), possibly because in the larger PheRS enzymes correct tRNA binding and orientation is dictated by multiple interactions across the fold, so that this role does not critically reside on the B3/B4 domain, which instead might have evolved to optimize its interactions with other domains within the enzyme. Taken together, these findings suggest that indels have directed the evolution of substrate binding in this class of enzymes.

**Fig 3.**
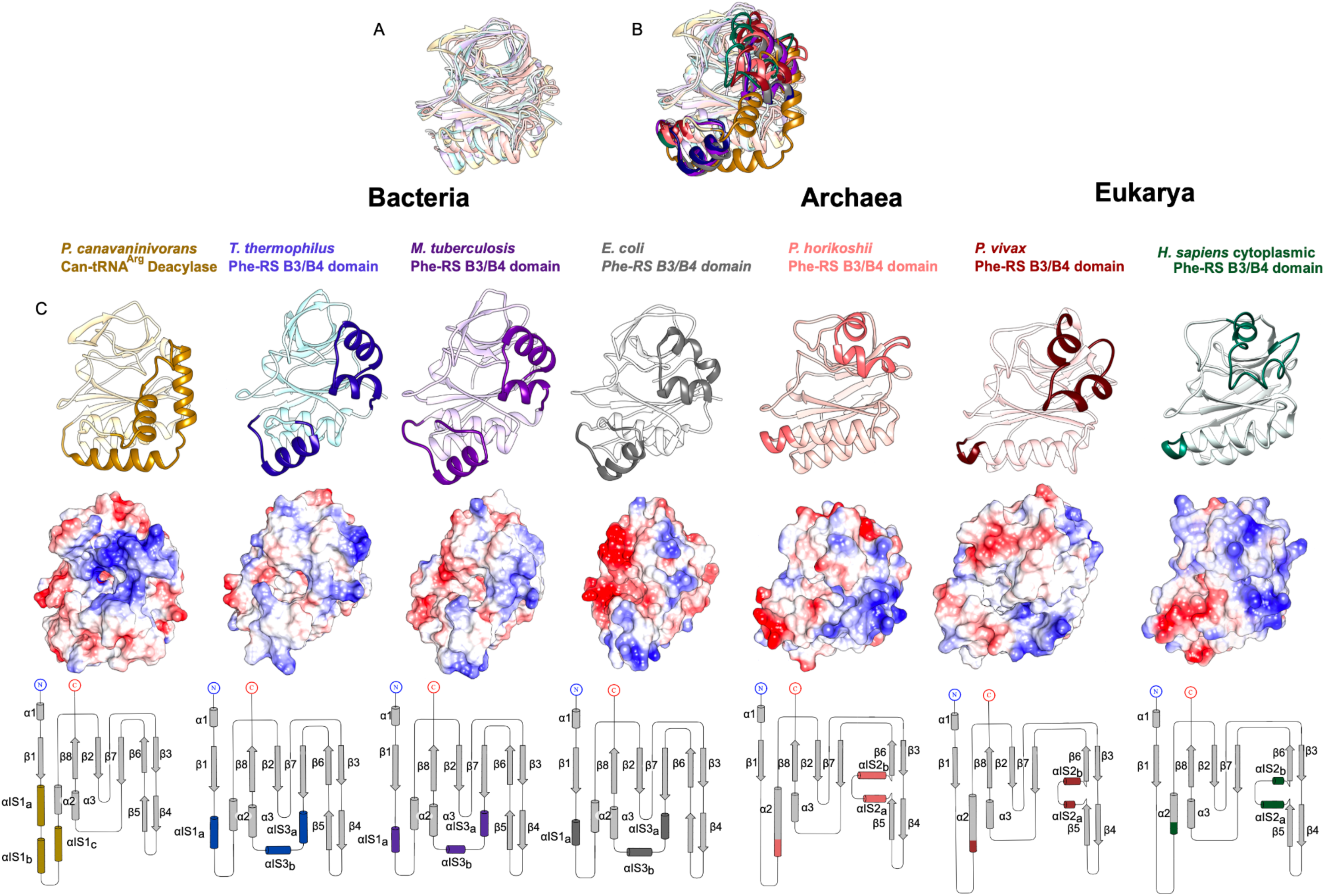
Structural comparison of CtdA and B3/B4 domains from PheRS proteins. CtdA and B3/B4 domains share a common fold core (**A.**) that carries species-specific indels (**B.**). The common core fold is represented in light colour shades and the indels are dark coloured, solid ribbons; **C.** Comparative overview of CtdA and B3/B4 domains of known structure shown in the same orientation. A selected set of representatives that show conserved features within their respective group is shown here for simplicity. For each structure, a ribbon representation (*upper*), a surface representation coloured according to electrostatic potential (red: -10 kcal/mol*e, blue: 10 kcal/mol*e) (*middle*), and a topology diagram (*bottom*) are shown. Colour-coding throughout is as in B., except for topology diagrams, where to ease comparative visualization the common core is depicted in grey and shown using a constant representation across scaffolds. Secondary structure elements are named according to their structural type and numerical order along the conserved fold or using the prefix IS if they belong to insertion sequences specific to enzyme subclass. For PheRS structures, the residue ranges displayed are *P. vivax* (PDB: 7BY6; residues 96-291), *T. thermophilous* (PDB: 2IY5; residues 201-397), *M. tuberculosis* (PDB: 7DB7; residues 211-412), *H. sapiens* (PDB: 3L4G; residues 103-291), *E. coli* (PDB: 3PCO; residues 201-400), *P. horikoshii* (PDB: 2CXI; residues 87-263).

**Fig 4:**
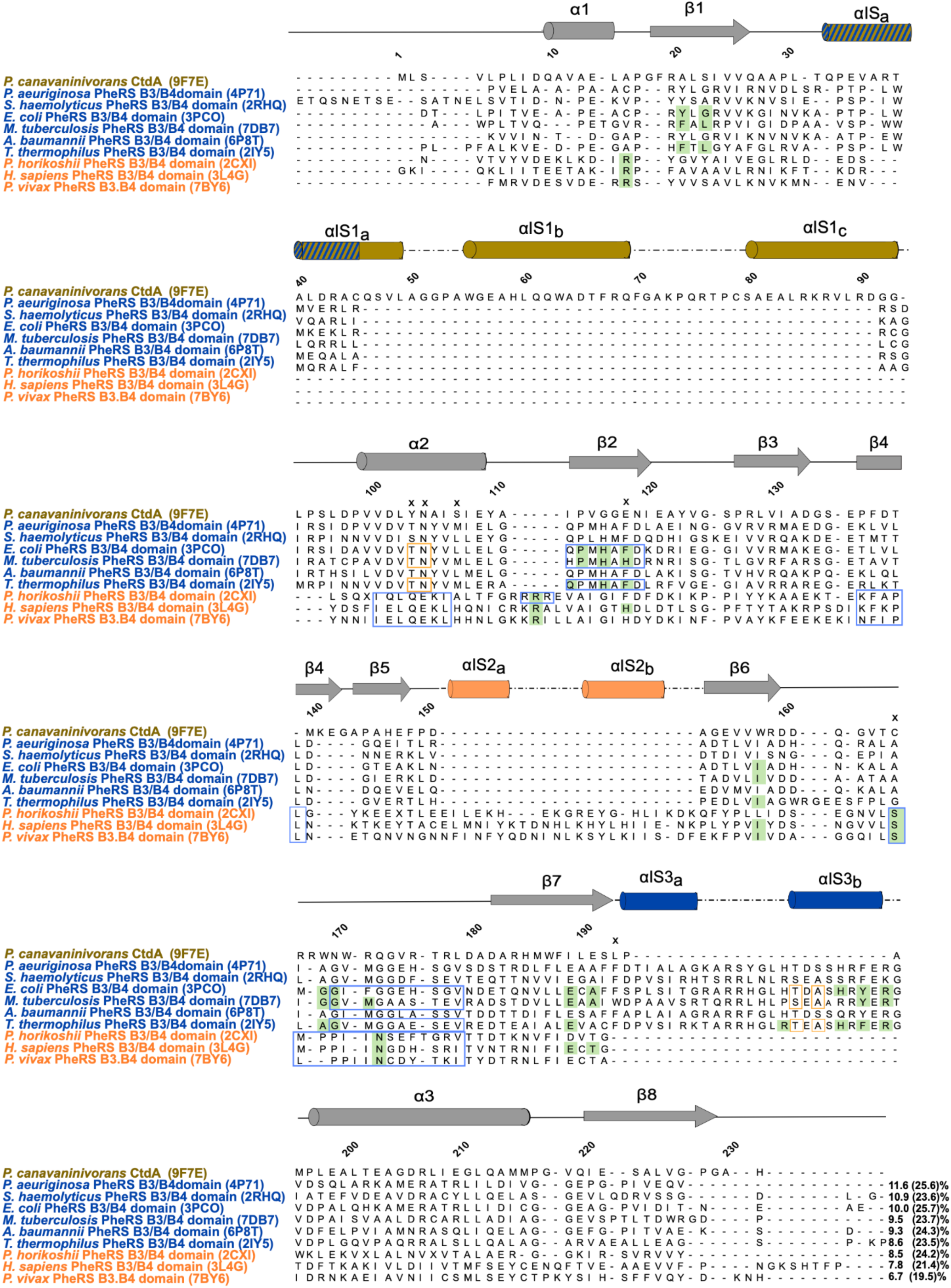
Structure-based sequence alignment of CtdA and B3/B4 domains from PheRS proteins of known 3D-structure. The 3D-structures of B3/B4 domains from PheRS proteins were obtained from the PDB entries with entry code indicated in brackets. Structures were aligned using VMD (Humphrey et al., 1996) to generate a structure-based alignment. Sequence identities and similarities (in brackets) were calculated using the Sequence Manipulation Suite (Stothard, 2000, www.bioinformatics.org/sms2/ident_sim.html) and are quoted (both as a percentage) at the end of each sequence. In the alignment, amino acid residues boxed in blue depict motifs that have been suggested to be involved in editing. Golden boxes indicate amino acids thought to position water molecules involved in substrate coordination. Residues highlighted in green interact with the aminoacyl substrate. Amino acids marked with an “X” are those of CtdA from *P. canavaninivorans* that have been mutated in the current study. Annotations for the individual sequences are based on data reported as follows: *Thermus thermophilus* (Moor et al., 2006), *Mycobacterium tuberculosis* (Wang et al., 2021), *Eschericha coli* (Mermershtain et al., 2011), *Plasmodium vivax* (Sharma et al., 2021), *Pyrococcus horikoshii* (Sasaki et al., 2006), and *Homo sapiens* PheRS (Finarov et al., 2010) *Pseudomonas aeuriginosa* PheRS (Abibi et al., 2014), *Acinetobacter baumannii* ( Baidin et al., 2021), *Staphylococcus haemolyticus* ( Evdokimov et al., 2008).

### Conservation of catalytic features is restricted to asparagine 105 in the oxyanion hole

Finally, we asked whether CtdA might share a general catalytic mechanism with B3/B4 domains from PheRS enzymes. To address this question, we performed a structure-based sequence alignment of CtdA and the B3/B4 domains listed above and studied sequence conservation to reveal potentially shared catalysis residues (**Fig 4**). We found that despite their close fold conservation, CtdA and B3/B4 domains share very low levels of sequence identity (ranging from 11.6% to 6.7%) or similarity (25.6-19.5%) (**Fig 4**). There is no discernible conservation of catalytic motifs across all groups, including residue candidates for coordinating the oxyanion and positioning the ribose ester to be cleaved. Thus, it cannot be established whether editing enzymes of this class share the same catalytic mechanism. Interestingly, CtdA is more similar to bacterial B3/B4 domains, with which it shares the strictly conserved residue N105 (CtdA numbering). In the *E. coli* PheRS editing site, this residue -and the preceding threonine rest-was proposed to coordinate the nucleophilic water molecule that attacks the substrate (Ling et al., 2007, Mermershtain et al. 2011). Our mutagenesis data indicate that residues Y104 and N105 in CtdA are also catalytically important (**Fig 2B**) and the crystal structure shows that these residues, contributed by helix α2 from the shared core fold, likely localize to the proximity of the canavanyl-ribose junction. Thus, it could be speculated that the asparagine residue is part of a shared catalytic mechanism across CtdA and bacterial B3/B4 editing domains. Conserved catalytic features beyond that residue are not observed. In the B3/B4 domain from *T. thermophilus,* residue E334 has been attributed a crucial role in the recognition of the Tyr aminoacyl moiety (Kotik-Kogan et al., 2005), while in the B3/B4 domain from *E.coli* three residues H265A and E334 have been described to be essential for catalysis as their mutation reduced editing activity (Ling et al., 2007). None of these residues, or others considered to be functionally relevant (annotated in **Fig 4**) are conserved in CtdA. They are also largely not conserved across B3/B4 domains of PheRS enzymes from bacterial and archeal/eukaryotic origin (**Fig 4**). Thus, further studies will be required to conclude on whether this enzyme class shares a catalytic mechanism.

### Conclusion

In this study, we set out to characterize CtdA, a canavanyl-tRNA^Arg^ deacetylase that constitutes the first example of a standalone, trans-editing enzyme homologous to the *cis*-editing B3/B4 domains of PheRS enzymes (Jani & Pappachan, 2022). Moreover, CtdA is the first trans-editing enzyme identified to act on a non-proteinogenic substrate, the arginine-like canavanine. Its relation to the editing domains of PheRS enzymes, which acts on tyrosyl-/phenyl-rests, is thus unexpected. The elucidation of the crystal structure of CtdA reveals that CdtA and B3/B4 domains from bacterial and archeal/eukaryotic origin constitute three subclasses of a same conserved 3D-fold that differ in the presence of type-specific indels. Even though sequence conservation across the subtypes is very low (**Fig 4**), the conserved part of the 3D-fold is remarkably similar (**Fig 3**). In all cases, a cavity between domains B3 and B4 (or the corresponding parts in CtdA) forms the binding site for the aminoacyl rest. The entrance to this cavity is shaped by an α-hairpin formation originating from the indel sequences, which although contributed by different parts of the protein chain localize to the same locus in the 3D-fold. In CtdA, this α-hairpin -part of the single, long indel IS1-contains positively charged residues that could contribute to the binding and recognition of the tRNA stem.

Despite their structural similarity, no conserved residues across these three subclasses of editing enzymes can be identified that suggest the same catalytic mechanism. For bacterial PheRS and archaeal ThrRS cis-editing domains, the editing mechanism has been proposed to follow substrate-assisted catalysis (Ling et al., 2007, Hussain et al., 2010). There, either the 2’OH or the 3’OH of the tRNAs A76 base is involved in the activation of a catalytic water molecule that is positioned within the editing site, which in turn makes a nucleophilic attack on the carbonyl carbon of the substrate. The function of those editing domains, therefore, appears to be the acceleration of the spontaneous hydrolysis of aa-tRNA by several orders of magnitude. In CtdA, a similar mechanism seems possible. *In silico* modelling of canavanine binding into the active site suggests that canavanine can be accommodated in the predicted binding pocket. Mutagenesis of two negatively charged residues at the deep end of the binding cavity (E118 and E191) showed both residues to be required for enzymatic activity. Further, we found residues Y104 and N105, which predictably are in proximity of the canavanyl-ribose junction, to be both also catalytically important. This finding is in agreement with previous reports on the B3/B4 domain of *E. coli* PheRS that proposed the two equivalent residues (T, N) to coordinate the nucleophilic water molecule that attacks the substrate (Ling et al., 2007, Mermershtain et al. 2011). The strict conservation of the asparagine residue across bacterial B3/B4 domains and CtdA (**Fig 4**) supports its importance for editing catalysis and hints at an ancestral relation of CtdA and the bacterial subclass of B3/B4 editing domains, which is supported by the closer sequence and structural relation between these two groups (**Fig 3 & Fig 4**). Furthermore, in this study we identify a number of enzymes that share high sequence conservation with CtdA, all being of bacterial origin and pointing to the potential spread of CtdA across bacteria in canavanine-rich habitats. Unfortunately, we were not able to obtain co-structures of CtdA in complex with substrates and, thus, we could not clarify the structural basis for the high degree of canavanine/arginine substrate discrimination observed in CtdA. The pKa values of the guanidino groups of canavanine and arginine differ drastically (7.0 vs 13.8, respectively; Boyar et al., 1982; Fitch et al., 2015). Thus, in addition to the 5-oxa-substitution, the changed protonation state of the guanidine group in these compounds could also serve CtdA as a means to differentiate between canavanine-and arginine-loaded tRNAs. Future studies will be required to reveal the molecular basis of substrate discrimination and editing activity by CtdA.

## Methods

### CtdA overexpression and purification

CtdA from *Pseudomonas canavaninivorans* (NCBI MBJ2347151.1) was overexpressed and purified as described previously (Hauth et al., 2023). In brief, expression plasmids were transformed into *E. coli* BL21(DE3) strain Gold (Agilent) and cultures were grown in Luria-Bertani medium supplemented with kanamycin (30 µg/ml) at 37°C up to an OD_600_ of 0.5. Protein expression was induced with 0.5 mM IPTG (isopropyl-β-D-thiogalactopyranosid) and cultures further grown for 16h at 18°C. Cells were harvested by centrifugation and lysed by sonication in 50 mM TrisHCl pH 8.0, 100 mM NaCl, 20 mM imidazole, protease inhibitor (cOmplete™ Mini, EDTA-free, Merck), 0.02 mg/ml lysozyme. Cell debris was removed by centrifugation. Protein purification was performed by immobilized metal affinity chromatography (IMAC) using Ni^2+^-NTA agarose. The His_6_-tag was removed by cleavage with TEV (Tobacco-Etch-Virus) protease and the sample was buffer exchanged into 50 mM TrisHCl pH 8.0, 100 mM NaCl and further purified by subtractive IMAC and size exclusion chromatography on a Superdex S75 16/60 column (GE Healthcare). The resulting protein samples were analysed by SDS-PAGE to ensure protein integrity and purity. Finally, the sample was concentrated to 17 mg/mL and immediately used in crystallization trials.

### AlphaFold

To assist phasing in crystallographic studies, a 3D-model of CdtA was calculated using AlphaFold v2.1.1 (Jumper et al., 2021). Calculations were executed locally at the University of Konstanz on a customized computer server consisting of a processor AMD Ryzen Threadripper Pro 3975 wx equipped with graphics card Nvidia GForce RTX 3090 and 256GB RAM running the Ubuntu 20.04.4 operative system. The template database included entries at the Protein Data Bank up to 22-12-2021. Six models were calculated and the relaxed model 0 was chosen as the molecular replacement search model.

### X-ray Crystallography

For crystallization, CtdA was concentrated to 23 mg/mL and incubated with canavanine at 1:2 molar ratio in 50 mM Tris pH 8.0, 100 mM NaCl. Crystals grew from solutions consisting of 0.6M K_3_PO_4_, 1.3M Na_2_HPO_4_, 0.2M Li_2_SO_4_, 0.1 M CAPS pH 10.5, 27% [w/v] PEG 8K, 0.2M AmSO_4_ in 96-well Intelli-Plates (Art Robbins) using the sitting-drop vapour-diffusion method at 18°C. Crystallization drops consisted of a 1:1 or 1:3 ratio of protein solution and reservoir solution and had a volume of 200:200 nl, 100:300 nl. For X-ray irradiation, crystals were cryoprotected with Paratone-N (Hampton Research) prior to flash-vitrification in liquid nitrogen.

X-ray diffraction data were collected on beamline PX1 at the Swiss Light Source synchrotron (Villigen,CH) under cryo-conditions. Diffraction data were processed using the XDS/XSCALE suite (Kabsch 2010). For phasing, the AlphaFold model of CdtA was used as a search model in molecular replacement, which was performed in Phenix.phaser (Adams et al., 2010). Manual model building was performed in Coot (Emsley et al., 2010) and model refinement was in phenix.refine (Liebschner et al., 2019) using anisotropic *B* factors and non-crystallographic symmetry (NCS) restraints. In the last refinement cycle, hydrogen atoms were added. The quality of the final model was assessed using MolProbity (Williams et al., 2018). No electron density was identified in the final model that could correspond to the canavanine compound. X-ray data statistics and model parameters are given in **Table 1**. Molecular images in this manuscript were produced using UCSF Chimera (Pettersen et al., 2004).

### *In silico* modelling of canavanine docked to CtdA

The structure of CtdA was parametrized using Chimera UCSF Dock Prep, which added hydrogens and charges calculated using ANTECHAMBER (Maier et al., 2015) applying the AMBER ff14SB force field for standard residues (Wang et al., 2006) and Gasteiger for other residues (Wang et al., 2004) with a box size of 39.5Å x 38.7Å x 42.9Å. Parametrized canavanine (entry ZINC3869452 in the compound database ZINC15; https://zinc.docking.org) was docked into CtdA using AutoDock Vina 1.1.2 (Trott et al., 2010). The merging of nonpolar hydrogen atoms and electron lone pairs was set to true. Waters and non-standard residues were included in the docking parameters. Hydrogen bonds were considered. Nine binding modes were tested, with an exhaustiveness of eight and a maximum energy difference of 3 kcal mol^−1^.

### Effect of point mutations on editing activity

CtdA variants were generated by whole plasmid overhang PCR using the wild-type CtdA overexpression plasmid as template. PCR was followed by Quick Ligation (NEB) and transformation by electroporation. The resulting clones were verified by sequencing (GATC, Eurofins) and the corresponding protein variants were purified as described above.

To determine the editing activity of the mutated protein variants, they were incubated with radioactively labelled tRNA as described before (Hauth et al., 2023). In summary, tRNA^Arg^ was *in vitro* transcribed using T7 RNA polymerase and purified by PAGE. tRNA was visualized by UV shadowing, excised and recovered from cut gel pieces using the crush and soak method and ethanol precipitation. Then, the purified tRNA was labelled with α-32P-ATP (Hartmann Analytic) using purified *E.coli* cca tRNA nucleotidyltransferase (ASKA collection, JW3028). Labelled tRNA was again purified by PAGE. Prior to the aminoacylation reaction, unlabelled tRNA was spiked with freshly prepared 32P labelled tRNA and refolded by heating to 98°C and slow cooling. Aminoacylation was performed using the ArgRS enzyme from *Pseudomonas canavaninivorans* (MBJ2348292.1) and either arginine or canavanine as a substrate. The reaction mixture consisted of ∼3 µM tRNA, 3 mM ATP, 0.0026 U/µl PPase, 0.13 µg/µl bovine serum albumin, 0.1 mM DTT, 6 µM ArgRS enzyme, 2 mM substrate, all in reaction buffer (50 mM HEPES, 25 mM KCl, 15 mM MgCl_2_, pH 7.5). The reaction was incubated for 2 hrs at 37°C after which it was quenched by the addition of 0.1 volumes of quenching buffer (1.2 M NaOAc, 0.1% [w/v] SDS, pH 4.0). Aminoacylated tRNA was precipitated using ice-cold ethanol and incubated for 1 hr at -80°C. Precipitated aminoacylated tRNA was pelleted by centrifugation and washed twice with 70% ethanol. Then, the tRNA was resuspended in the reaction buffer and incubated with 1 µM CtdA protein or its respective variants for 10 min at room temperature. The reaction was then quenched again using quenching buffer and the tRNA digested using P1 nuclease for 1 hr at room temperature. Following, the digested samples were separated by thin layer chromatography on polyethylenimine cellulose F plates (Supelco, Merck). Running buffer consisted of 100 mM ammonium acetate in 5% [v/v] acetic acid. Plates were dried on air and radiographs recorded on a phosphorimager (GE Healthcare Life Science). ImageJ (1.53t, NIH, USA) was used to quantify and evaluate signal intensities.

## Data availability

Structure coordinates and experimental diffraction data have been deposited with the PDB (entry 9F7E). X-ray diffraction images have been deposited with Zenodo (http://doi.org/10.5281/zenodo.11109289)

## Acknowledgments

We gratefully acknowledge financial support from the European Research Council [ERC CoG 681777 ‘RiboDisc’ to J.S.H.].

## Author contribution

FH, JF, JH, OM conceived the study; FH, JF performed cloning, protein production, and crystallization; JF, NT, OM performed crystallographic structural elucidation; JF, NT, OM, FH performed bioinformatics analysis; OM, FH, NT wrote the manuscript and all authors made revisions.

## SUPPLEMENTARY MATERIALS

**Fig S1:**
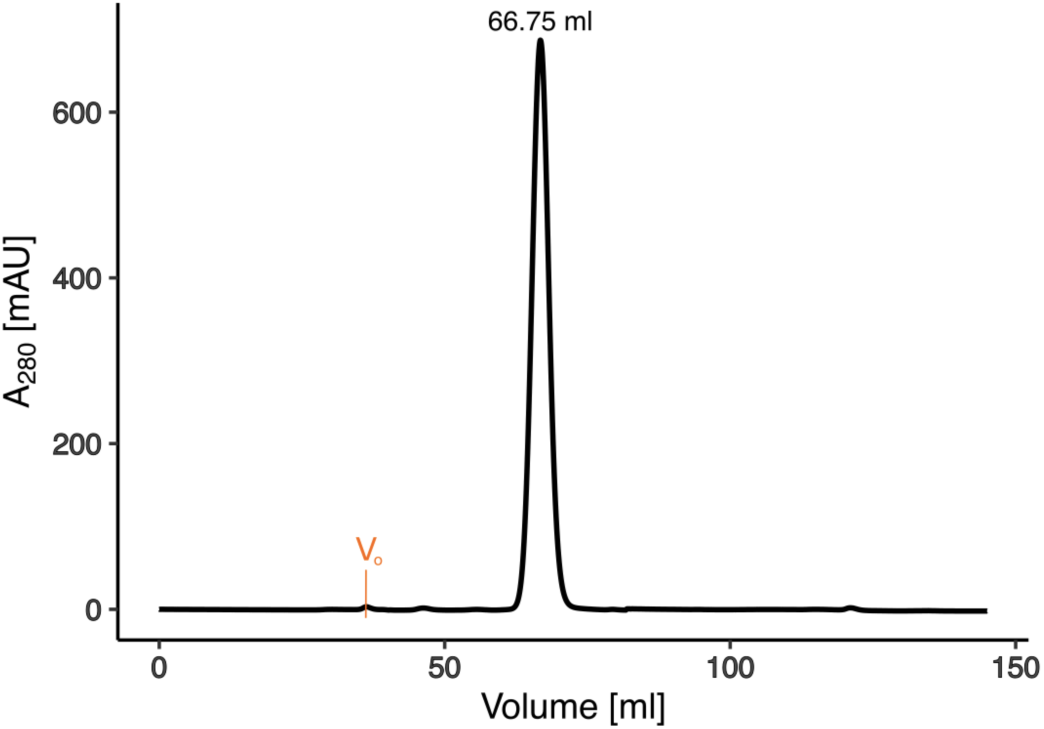
Size exclusion chromatogram of CtdA. The CtdA sample was loaded onto a Superdex 75 16/600 column (Cytiva) in 50mM Tris, 100mM NaCl, pH 8.0. A single, symmetric peak was observed at an exclusion volume of 66.75 ml, which corresponds to a molecular mass of 25.2 kDa. The standard curve was determined using the Gel Filtration Cal Kit Low Molecular Weight (Cytiva). The column void volume is shown in orange.

**Fig S2:**
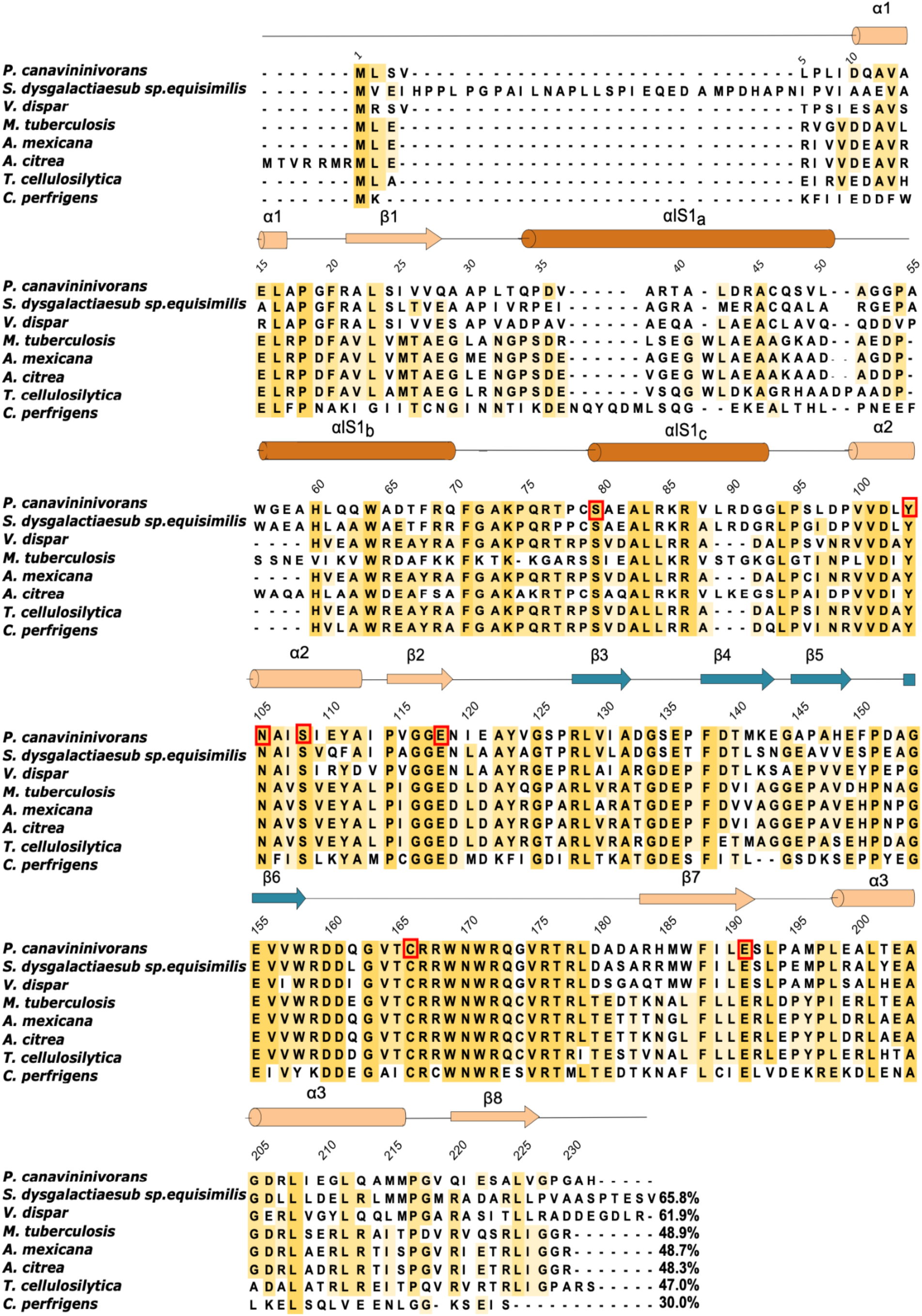
Alignment of sequences closely-related to CtdA. Sequence alignment of CtdA from *P. canavaninivorans* with closely related sequences of putative canavanyl-tRNA^Arg^ deacylases from *S. dysgalactiaesub sp equisimilis* (VTS39953.1) *V. dispar* (MBS6383566.1) , *M. tuberculosis* (CNF07723.1), *A. mexicana* (SNR39375.1), *A. citrea* (NYE09768.1), *T. cellulosilytica* (WP_182704228.1), and *C. perfrigens* (WP_004456252.1) . Orange shadowing highlights sequence conservation. The numbering is given for the sequence of CtdA from *P. canavaninivorans.* The secondary structure elements of *P. canavaninivorans* CtdA are depicted on top of the alignment and are as revealed by the crystal structure. Secondary structure elements are colored according to homology to the B3/B4 domains of PheRA and match the colouring scheme of Fig 1 in the main text; namely, homology to domain B3 is shown in gold and to B4 in blue. Dark gold marks the long indel sequence within domain B3 that is specific to CtdA enzymes. Residues mutated in this study are boxed in red. Sequence identities were calculated in Jalview v2.11.3.2. (https://www.jalview.org; Waterhouse et al., 2009)

**Fig S3.**
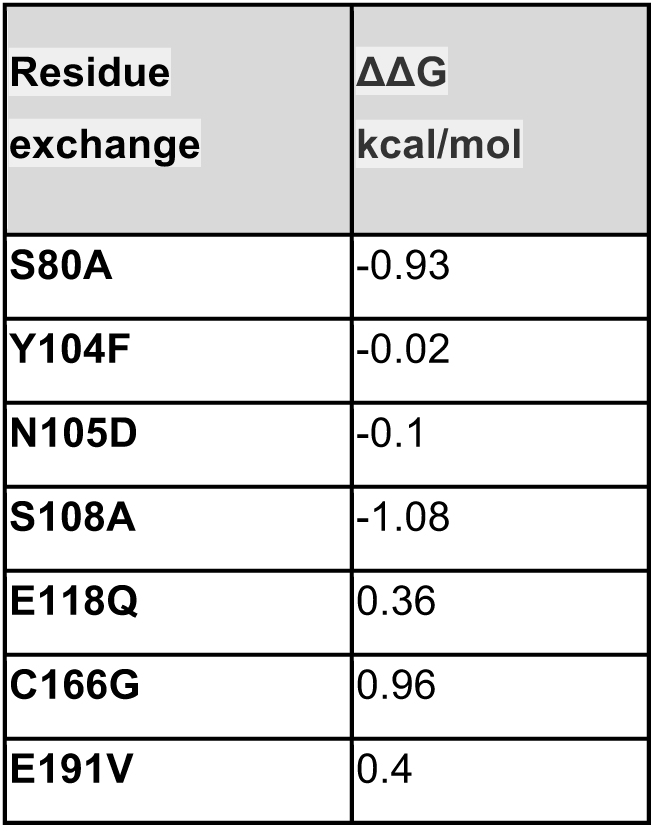
*In silico* mutation stability check of CtdA point mutants. FoldX v.5 (https://foldxsuite.crg.eu; Schymkowitz et al., 2005) was used to calculate positional destabilization effects in point mutational screens. For this, the crystal structure of CtdA was energy minimised in FoldX and then specific residues mutated *in silico* in turn. Default settings were used for the program. For each mutated residue, FoldX calculates the difference in Gibbs free energy of folding between mutant and wild-type form of the protein (ΔΔG = ΔGmutant – ΔGwt). While there is no established ΔΔG threshold value to classify instability in FoldX, typically variations within 1 kcal/mol are considered well tolerated by the protein structure and not to alter stability noticeably.

**Fig S4:**
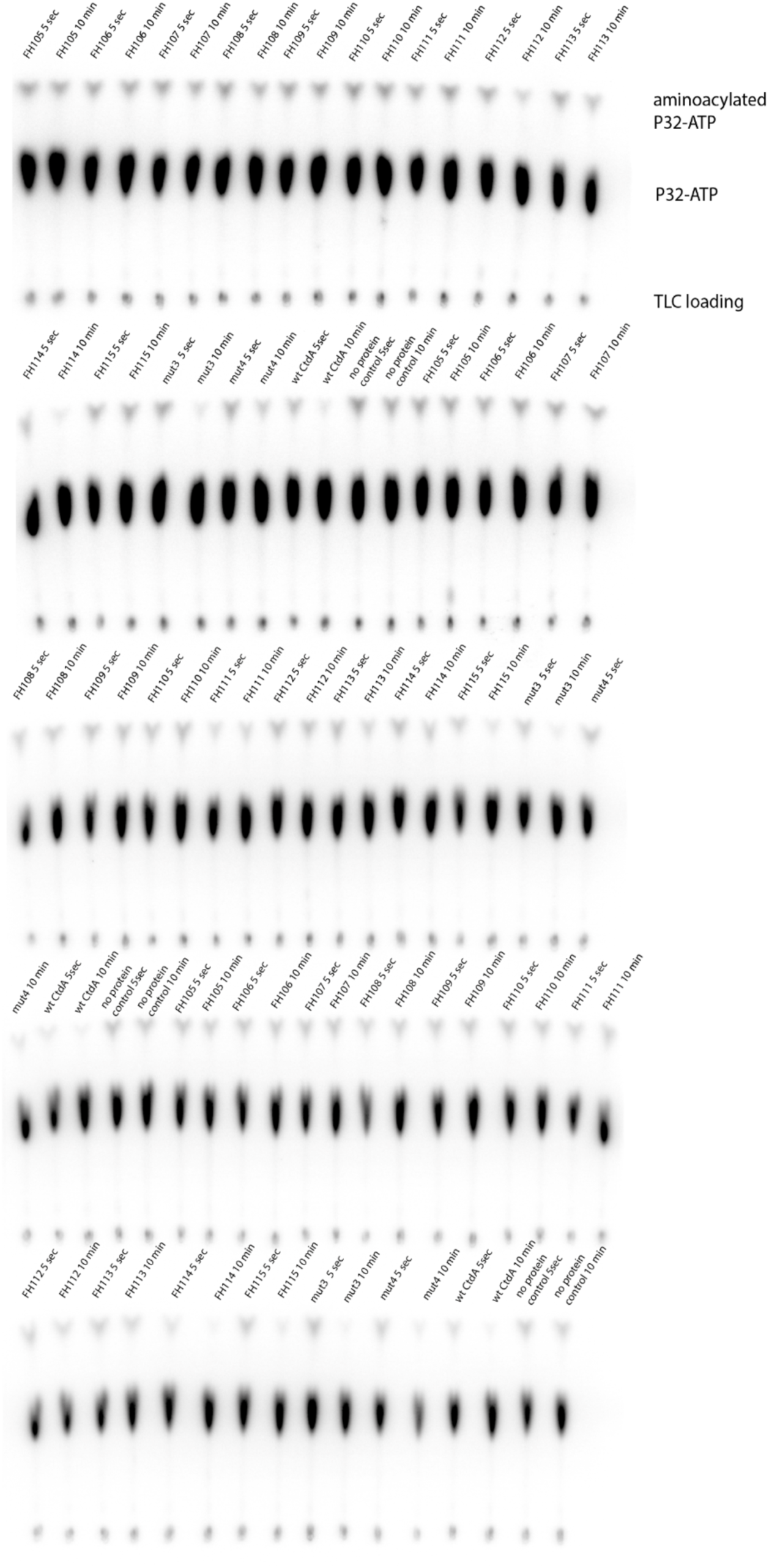
Thin-layer-chromatography radiographs used for activity assessment of CtdA and its mutants. Samples were quenched 5 secs or 10 min after the addition of the respective protein, digested by RNAse treatment and subjected to TLC. Radiographs were evaluated the following day using ImageJ by comparing the ratio between the aminoacylated and the non-aminoacylated peak. mt3: S80A, mt4: S108A, FH106: E191V, FH108: Y104F, FH110: N105D, FH112, E118Q, FH115: C166G

